# Neural regulation of infection resolution prevents ER stress and tissue damage

**DOI:** 10.64898/2026.05.08.723879

**Authors:** Phillip Wibisono, Noor Hamoud, Sagi Levy, Jingru Sun

## Abstract

Neural regulation of immunity is increasingly recognized, yet how the nervous system promotes infection resolution after pathogen clearance remains poorly understood. Using *Caenorhabditis elegans*, we identify the AIA interneurons as key regulators of both infection and recovery. Acute silencing or genetic ablation of AIA neurons during *Salmonella enterica* infection reduces host survival, causes excessive activation of conserved immune and stress pathways, including PMK-1/p38 MAPK, insulin/IGF-1 signaling, and the XBP-1-mediated unfolded protein response (UPR), and exacerbates intestinal tissue damage. Strikingly, selective silencing of AIA neurons after pathogen clearance markedly impairs recovery, demonstrating that neural activity is required not only for host defense but also for infection resolution. This defect is rescued by *xbp-1*, but not *pmk-1* or *daf-16*, knockdown, identifying unresolved ER stress as the principal driver of post-infection mortality. Consistently, AIA silencing during recovery sustains UPR activation and worsens epithelial barrier damage. Together, our findings establish AIA interneurons as central regulators of immune homeostasis that promote infection resolution by limiting excessive ER stress and preserving tissue integrity.

## Introduction

Increasing evidence indicates that upon pathogen infection, the host nervous and immune systems interact to generate coordinated protective responses (1, 2). In mammals, neural regulation of immunity occurs through hormonal, autonomic, and neuropeptidergic pathways that integrate inflammatory responses with environmental cues and physiological state (3–5). Despite these advances, many mechanistic details of neuroimmune communication remain poorly understood, in part because the complexity of mammalian nervous and immune systems poses significant experimental challenges. In recent years, the nematode *Caenorhabditis elegans* has emerged as a powerful model organism for studying neuroimmune regulation, owing to its simple, well-defined nervous system and an immune system that resembles mammalian innate immunity in several key respects (6). Although *C. elegans* lacks professional immune cells, it mounts robust innate immune and stress responses through conserved pathways, including the PMK-1/p38 MAPK pathway, the insulin/IGF-1 signaling, and the unfolded protein response (UPR) (7). Studies using this model system have revealed unprecedented detail regarding the molecules, cells, and signaling pathways involved in neural regulation of immunity (6). For example, we have identified and characterized three distinct neuroimmune regulatory circuits (8–12), and others have described serotonergic, dopaminergic, cholinergic, and neuropeptidergic pathways that modulate immune responses (13–18).

A recurring theme emerging from these studies is that most neuroimmune regulatory circuits function to suppress, rather than activate, immune and stress responses. Inactivation of such neural circuits frequently results in increased expression of immune genes and enhanced survival against pathogen infection (6). This observation raises a fundamental question: why would neural circuits dampen immune responses that are clearly required for host defense? We propose that the primary functions of neuroimmune regulation are to prevent excessive activation of immune and stress responses during infection and to promote resolution of these responses once pathogens are cleared. While acute activation of immune and stress pathways is beneficial during infection, excessive or prolonged signaling can cause cellular dysfunction and pathology, leading to tissue damage and even host death (19, 20). After pathogen clearance, immune responses must be actively resolved. Failure to do so can result in uncontrolled inflammation, which is now recognized as a unifying mechanism underlying many inflammatory diseases, including arthritis, sepsis, and inflammatory bowel disease (21). Consistent with this idea, Serhan and colleagues showed that vagotomy, surgically removing the vagus nerve, delays resolution of *Escherichia coli* infection and peritonitis in mice, indicating that immune resolution is under neural control (22, 23). However, the precise mechanisms by which the nervous system regulates immune resolution remain unclear. Studies of defined neuroimmune regulatory circuits in *C. elegans* over the past decade have uniquely positioned this system to address two central questions: how neural circuits prevent overactivation of immune and stress responses during infection, and how they promote immune resolution during recovery.

Many sensory neurons (e.g., ASH, ASI, AWB, AWC, ASJ, ADF, ASG, CEP, AQR, PQR, and URX) contribute to *C. elegans* defense against pathogen infection, although the specific sensing functions of individual neurons in this context remain largely unknown (8, 12, 13, 15, 16, 18). Notably, most of these sensory neurons also regulate behavioral responses to environmental cues, including pathogen-associated signals, suggesting that the nervous system can detect pathogens either directly or indirectly (6). Because different sensory neurons likely respond to distinct pathogen-derived cues, studies focused on individual sensory neurons are unlikely to uncover general regulatory principles. In contrast, many pathogen-responsive sensory neurons converge onto a small number of first-layer interneurons, including AIA, AIB, AIZ, and AIY (24, 25). These interneurons integrate diverse sensory inputs and play central roles in decision-making processes that shape behavior and physiology (25). AIA and AIY receive inputs from chemosensory neurons involved in pathogen detection and food perception and regulate downstream circuits controlling locomotion and behavioral state (26, 27). Their position at the intersection of sensory processing and organism-wide output makes them strong candidates for coordinating immune homeostasis in response to infection.

Here, we investigate the roles of the AIA and AIY interneurons in regulating innate immune and stress responses during both active infection and post-infection recovery in *C. elegans*. Using acute neuronal silencing, genetic ablation, and recovery-specific assays, we show that AIA neuronal activity, but not AIY activity, restrains excessive activation of conserved immune and stress pathways during infection and promotes infection resolution by suppressing the XBP-1-mediated UPR. Loss of AIA activity results in persistent stress signaling, increased tissue damage, and reduced host survival. Together, these findings identify AIA interneurons as key regulators of immune homeostasis and infection resolution, revealing a critical role for interneurons in coordinating host defense and recovery following pathogenic challenge.

## Results

### Silencing AIA neurons reduces host survival during pathogen infection

Neuronal regulation of innate immune responses to pathogen infection is increasingly recognized, yet the precise mechanisms underlying such regulation remain unclear. To address this gap, we examined the roles of the AIA and AIY interneurons in *C. elegans* defense against *Salmonella enterica* SL1344 infection. These neurons were selected because they are first-layer interneurons that integrate sensory inputs and participate in decision-making processes, positioning them as strong candidates for modulating survival outcomes (24, 25).

To acutely inhibit neuronal activity, we employed transgenic strains expressing histamine-gated chloride channel 1 (HisCl) specifically in AIA neurons, AIY neurons, or both neuron types (i.e., *AIA::HisCl*, *AIY::HisCl*, and *AIA+AIY::HisCl* animals). Exposing these HisCl animals to histamine silences the target neurons within minutes, and neuronal inactivation is stable during histamine exposure but reversible with histamine removal (28, 29). Because *C. elegans* lacks endogenous histamine receptors, exogenous histamine is physiologically inert in the nematode and does not alter its biology (28, 29). Consistent with this, histamine treatment alone had no effect on the lifespan of wild-type (WT) animals or on their survival following *S. enterica* infection (Figures 1A and 1B). Moreover, histamine exposure did not alter infection-induced expression of typical genes in conserved immune and stress response pathways, including the PMK-1 pathway, the insulin/IGF-1 signaling, and the XBP-1-mediated UPR (Figure 1C) (9, 30, 31). These results indicate that histamine treatment does not affect *S. enterica* virulence or *C. elegans* immune signaling on its own. In the absence of histamine, survival following *S. enterica* infection was comparable among WT, *AIA::HisCl*, *AIY::HisCl*, and *AIA+AIY::HisCl* animals (Figure 1D), confirming that HisCl expression itself does not influence host survival. In contrast, histamine-induced silencing of AIA neurons, either alone or together with AIY neurons, resulted in a significant reduction in survival in *AIA::HisCl* and *AIA+AIY::HisCl* animals relative to WT controls (Figure 1E), suggesting that AIA neuronal activity contributes to host survival during the infection. By contrast, silencing AIY neurons alone did not significantly affect survival (Figure 1E), indicating that AIY activity is not essential for survival under these conditions and that the protective effect is specifically mediated by AIA neurons rather than a general property of interneuron function.

**Figure 1.**
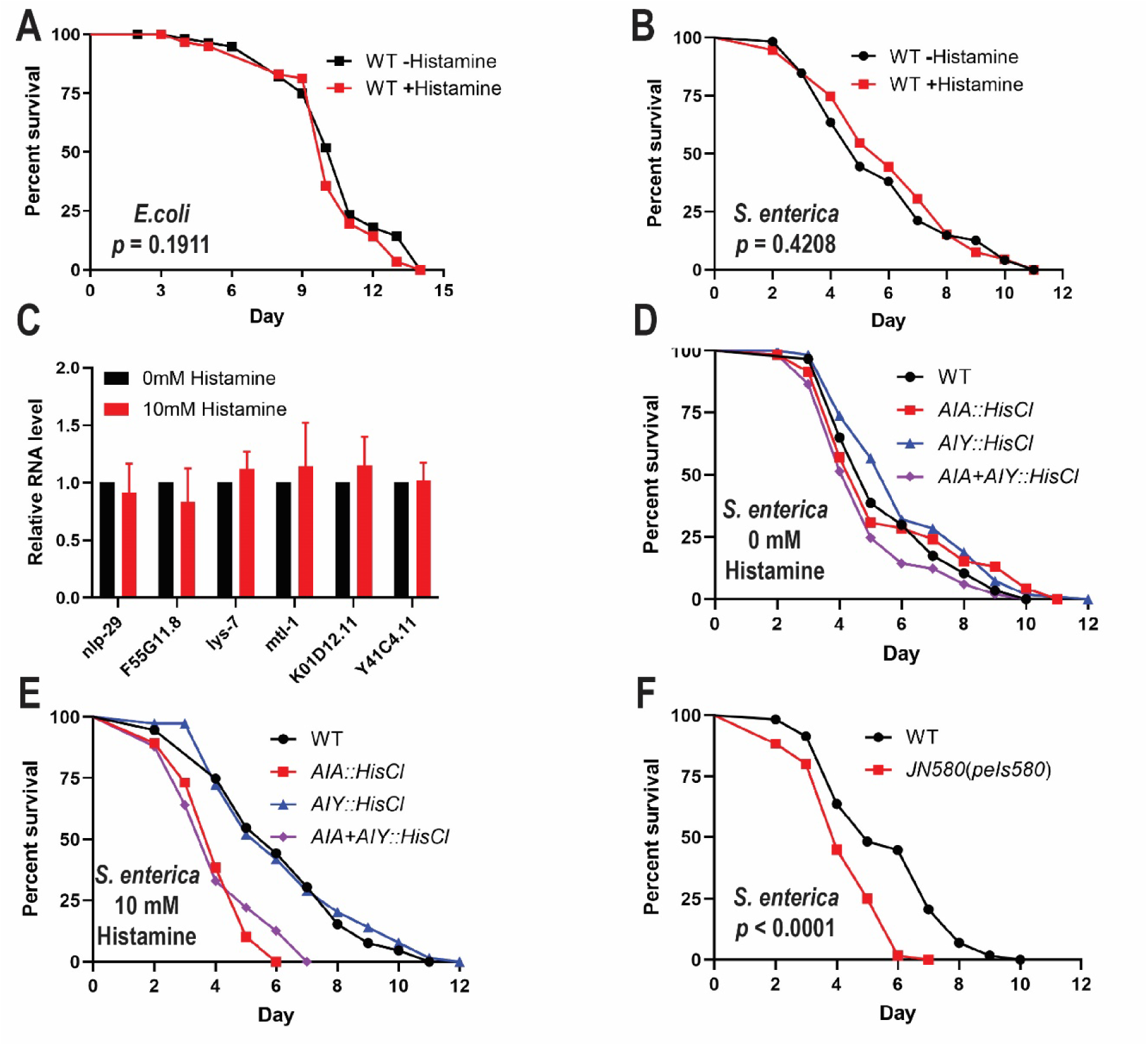
Silencing AIA neurons reduces host survival during pathogen infection. **(A)** WT animals were grown in the presence or absence of 10 mM histamine and scored for survival over time. **(B)** WT animals were grown in the presence or absence of 10 mM histamine, exposed to *S. enterica*, and scored for survival over time. **(C)** WT animals were grown in the presence or absence of 10 mM histamine and exposed to *S. enterica*. qRT-PCR was performed to measure the expression levels of typical genes in the PMK-1 immune signaling pathway (*nlp-29* and *F55G11.8*), the insulin/IGF-1 pathway (*lys-7* and *mtl-1*), and the XBP-1 UPR pathway (*K01D12.11* and *Y41C4.11*). The graph is the combined results of three independent experiments. Error bars represent SEM. Values shown are normalized expression levels relative to animals grown with 0 mM histamine. **(D)** WT, *AIA::HisCl*, *AIY::HisCl*, and *AIA+AIY::HisCl* animals were exposed to *S. enterica* in the absence of histamine and scored for survival over time. **(E)** WT, *AIA::HisCl*, *AIY::HisCl*, and *AIA+AIY::HisCl* animals were exposed to *S. enterica* in the presence of 10mM of histamine and scored for survival over time. **(F)** WT and AIA-ablated animals *JN580(peIs580)* were exposed to *S. enterica* and scored for survival over time. For lifespan or survival assays, each graph is representative of three independent replicate experiments. Each experiment included *n* = 60 animals per strain. *p*-values represent the significance levels relative to controls. In (A), *p* = 0.1911. In (B), *p* = 0.4208. In (D), relative to WT: *AIA::HisCl*, *p* = 0.6522; *AIY::HisCl*, *p* = 0.1069; *AIA+AIY::HisCl*, *p* = 0.0727. In (E), relative to WT: *AIA::HisCl*, *p* < 0.0001; *AIY::HisCl*, *p* = 0.6369; *AIA+AIY::HisCl*, *p* < 0.0001. In (F), *p* < 0.001.

To validate the requirement for AIA neurons in host defense, we examined animals in which AIA neurons were genetically ablated (*JN580(pels580)* animals). Upon *S. enterica* infection, AIA-ablated animals exhibited markedly reduced survival compared to WT animals (Figure 1F), closely phenocopying histamine-treated *AIA::HisCl* animals. Together, these findings demonstrate that AIA neurons play a critical role in promoting host survival during pathogen infection.

### Silencing AIA neurons causes excessive immune and stress responses to pathogen infection

While activation of innate immune and stress response pathways is essential for host defense against pathogen infection, excessive or prolonged activation can be detrimental, leading to tissue damage and reduced survival (32, 33). Because silencing AIA neurons reduced host survival (Figure 1, D and E), we hypothesized that AIA neurons help maintain a balanced innate immune and stress response to promote survival during infection.

To test this hypothesis, we examined the activity of key conserved immune and stress response pathways, including the PMK-1 pathway, the DAF-2/DAF-16 insulin signaling, and the XBP-1-mediated UPR, by measuring the expression of established pathway target genes. During *S. enterica* infection, *AIA::HisCl* and *AIA+AIY::HisCl* animals with histamine-silenced AIA neurons exhibited significantly elevated expression of most immune and stress response genes compared to WT animals (Figure 2). In contrast, gene expression levels in *AIY::HisCl* animals were comparable to those observed in WT animals (Figure 2). These results suggest that AIA activity is specifically required for suppressing gene overexpression of these immune and stress response pathways during infection, while AIY activity is not required.

**Figure 2.**
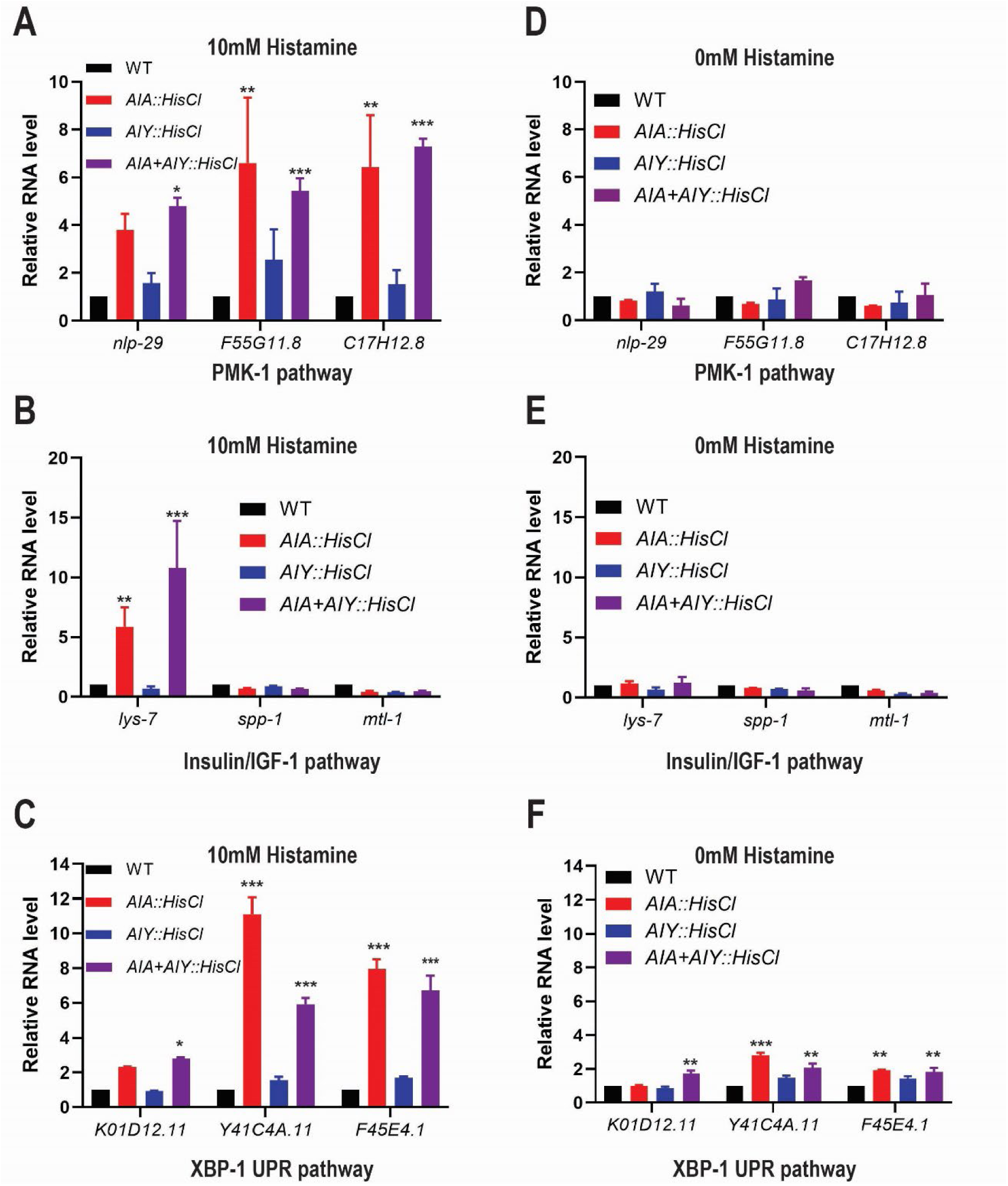
AIA neurons regulate gene expression in immune and stress response pathways during *S. enterica* infection. L4-stage WT, *AIA::HisCl*, *AIY::HisCl*, and *AIA+AIY::HisCl* animals were exposed to *S. enterica* for 4 hours in the presence (A-C) or absence (D-F) of 10mM histamine. RT-qPCR was performed to measure the mRNA levels of typical genes in the PMK-1 immune signaling pathway (**A, D**), the insulin/IGF-1 pathway (**B, E**), and the XBP-1-mediated UPR pathway (**C, F**). Bars represent the means ±SEM, *n* = 3 independent replicates. Asterisks (*, **, ***) denote significant difference (*p* < 0.05, 0.01, 0.001) relative to WT.

Notably, both *AIA::HisCl* and *AIA+AIY::HisCl* animals exhibited modestly elevated expression of XBP-1-regulated UPR genes relative to WT animals even in the absence of histamine treatment (Figure 2F). Consistent with previous reports of histamine-independent effects in some HisCl strains (29), the elevated basal UPR gene expression could reflect either strain-specific differences in gene expression or unintended basal silencing of AIA activity resulting from basal HisCl activity. To assess whether HisCl expression itself caused substantial impairment of AIA activity in the absence of histamine, we measured odor-evoked calcium responses in AIA neurons expressing both GCaMP and HisCl. AIA neurons exhibited robust responses across a 100-fold range of diacetyl concentrations, with larger responses observed at higher odor concentrations, indicating that AIA retains substantial sensory-evoked activity under these conditions (Figure S1). In contrast, histamine exposure strongly suppressed these odor-evoked responses, confirming efficient histamine-dependent silencing (Figure S1). Although these experiments do not exclude the possibility of subtle basal effects of HisCl expression on AIA excitability, they argue against substantial AIA silencing in the absence of histamine and support the use of histamine treatment as an effective acute manipulation of AIA activity. Thus, the elevated basal UPR gene expression observed in the absence of histamine is unlikely to be explained by substantial AIA silencing, and instead could reflect strain-specific differences in basal gene expression.

Overall, our findings indicate that AIA neurons act to restrain excessive activation of innate immune and ER stress responses during pathogen infection, and that loss of this regulation produces harmful overactivation that compromises host survival.

### AIA neurons promote infection resolution through regulation of the XBP-1-mediated UPR

Our infection-phase analyses showed that AIA neurons suppress excessive activation of innate immune and stress response pathways, raising the question of whether AIA neurons also contribute to the resolution of these responses after pathogen clearance. Effective recovery requires not only elimination of the pathogen but also timely attenuation of immune and stress signaling to restore tissue homeostasis (34). To directly assess the role of AIA neurons during the resolution phase, we employed a recovery assay adapted from Aballay and colleagues (35, 36) to isolate neuronal function specifically during recovery.

In this assay, synchronized WT and *AIA::HisCl* animals were exposed to *S. enterica* (or *E. coli* OP50 as a control) for 24 hours, washed extensively, and transferred to recovery plates seeded with *E. coli* OP50 supplemented with ampicillin to eliminate residual *S. enterica*. To selectively silence AIA neurons during recovery, animals were briefly pre-exposed to 10 mM histamine and maintained on recovery plates containing 10 mM histamine, whereas control animals were handled identically but without histamine treatment (Figure 3A). This experimental design enabled us to assess the contribution of AIA neuronal activity specifically during the recovery phase following infection.

**Figure 3.**
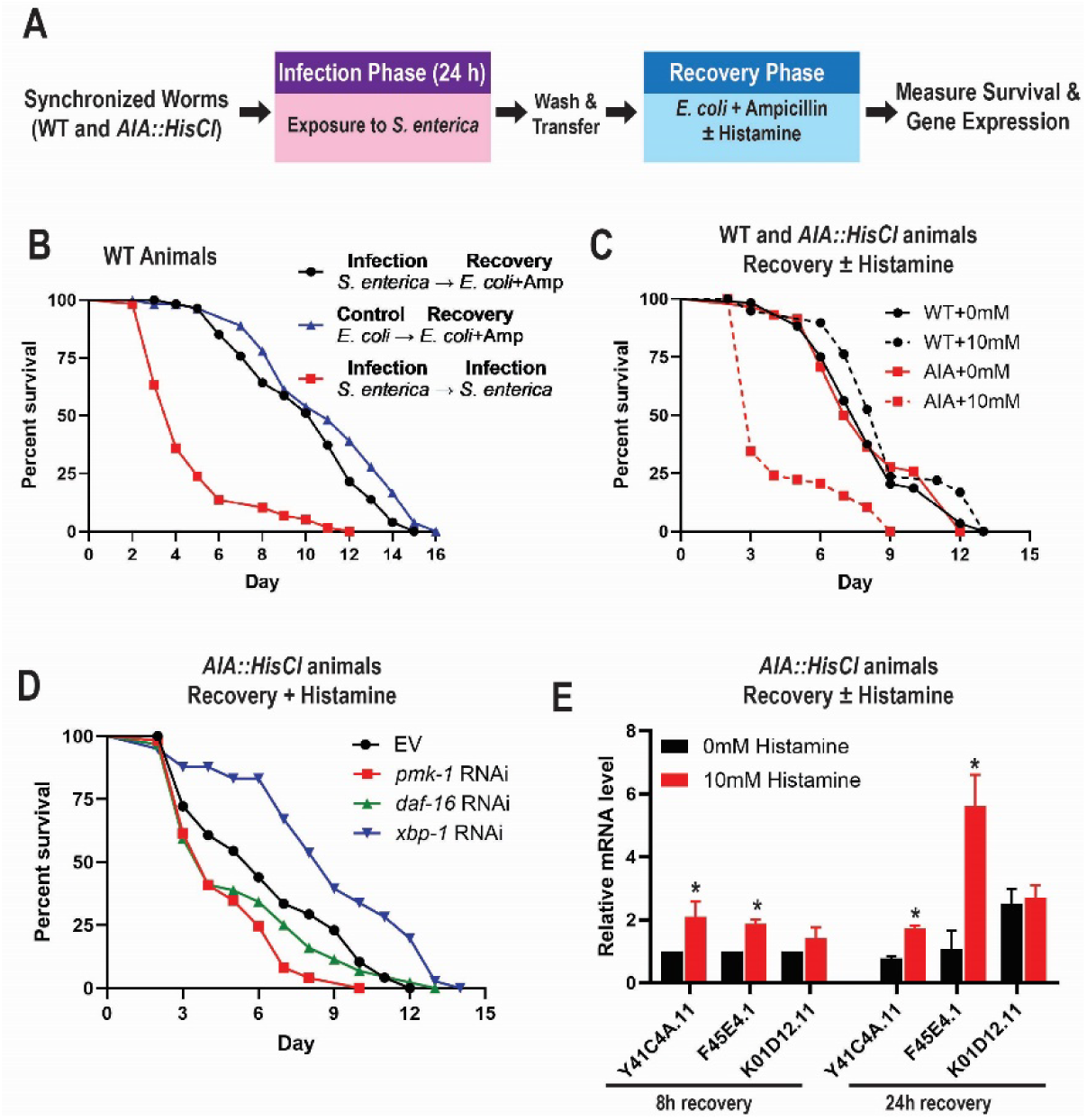
AIA neurons promote infection recovery by regulating the XBP-1-mediated UPR. **(A)** Schematic of the recovery assay. Synchronized WT and *AIA::HisCl* animals were exposed to *S. enterica* (or *E. coli* OP50 as a control) for 24 h, washed extensively, and transferred to recovery plates seeded with *E. coli* OP50 supplemented with ampicillin to eliminate residual *S. enterica*. To selectively silence AIA neurons during recovery, animals were briefly pre-exposed to 10 mM histamine and maintained on recovery plates containing 10 mM histamine. Control animals were handled identically but with 0 mM histamine treatment. **(B)** Synchronized 65-hour old WT animals were exposed to *E. coli* or *S. enterica* for 24 hours and then transferred to recovery plates containing either *E. coli* + ampicillin or *S. enterica*. Animals were scored for survival over time. The graph is representative of three independent experiments with *n* = 60 animals per strain. *p*-values represent the significance levels of survival relative to *E. coli*→*E. coli*+Amp: *S. enterica*→*E. coli*+Amp, *p* = 0.0319; *S. enterica*→*S. enterica*, *p* < 0.0001. **(C)** Synchronized 65-hour old WT and *AIA::HisCl* animals were exposed to *S. enterica* for 24 hours before being transferred to recovery plates seeded *E. coli* containing either 0mM or 10mM of histamine. Survival was monitored over time. The graph is representative of three independent experiments with *n* = 60 animals per strain. *p*-values represent the significance levels of survival relative to the WT + 0mM histamine: WT + 10mM histamine, *p* = 0.1642; *AIA::HisCl* + 0mM histamine, *p* = 0.3739; *AIA::HisCl* + 10mM histamine, *p* < 0.0001. **(D)** Synchronized 65-hour old *AIA::HisCl* animals fed dsRNA against specific genes or empty vector (control) were exposed to *S. enterica* for 24 hours before being transferred to recovery plates seed with *E. coli* and containing 10 mM histamine. Survival was scored over time. The graph is representative of three independent experiments with *n* = 60 animals per strain. *p*-values represent the significance levels of survival relative to EV: *pmk-1*, *p* = 0.0015; *daf-16, p* = 0.2167; *xbp-1, p* = 0.0001. **(E)** Synchronized 65-hour old *AIA::HisCl* animals were subjected to recovery assays in the presence or absence of 10mM histamine. qRT-PCR was performed to measure the expression levels of typical genes in the XBP-1 UPR pathway. The graph is the combined results of three independent experiments. Error bars represent SEM. Values shown are normalized expression levels relative to animals grown with 0 mM histamine at 8h recovery. Asterisks (*) denote significant difference (*p* < 0.05) between 0 mM and 10 mM histamine for individual genes at each recovery time point.

As shown in Figure 3B, animals infected with *S. enterica* and subsequently treated with ampicillin exhibited significantly improved survival compared with animals that remained continuously infected. Notably, survival of infected animals following ampicillin treatment closely approximated that of animals that were never exposed to *S. enterica* (Figure 3B). These results demonstrate that *S. enterica* infection can be effectively cleared by ampicillin treatment and establish this regimen as a tractable model for studying post-infection recovery, consistent with the tetracycline-based recovery model described by Aballay and colleagues (35, 36).

We applied this recovery paradigm to examine the role of AIA neurons during resolution. As expected, WT animals showed no significant difference in survival with or without histamine treatment, confirming that histamine itself does not affect recovery (Figure 3C). In contrast*, AIA::HisCl* animals exhibited a marked reduction in survival when AIA neurons were silenced with histamine during recovery compared with *AIA::HisCl* controls without histamine treatment (Figure 3C). These findings indicate that AIA neuronal activity is required during the recovery phase following *S. enterica* exposure to support host survival.

To identify the downstream pathways mediating this effect, we targeted key regulators of innate immunity and stress responses by RNAi in *AIA::HisCl* animals undergoing recovery. Knockdown of *xbp-1*, but not *pmk-1* or *daf-16*, significantly rescued the survival defect of AIA-silenced animals (Figure 3D), indicating that excessive activation of the XBP-1-mediated UPR is detrimental during recovery. Our qRT-PCR analysis confirmed that the expression of XBP-1 target genes in recovering *AIA::HisCl* animals were significantly higher in the presence of histamine than in its absence (Figure 3E). Together, these results demonstrate that AIA neurons play a critical role in infection resolution by restraining XBP-1-mediated UPR signaling during recovery. In combination with their role in limiting excessive immune and stress responses during active infection, these findings reveal a dual function for AIA neurons in coordinating defense and recovery programs to promote host survival following pathogenic challenge.

### AIA neuronal activity is required for limiting tissue damage during infection and recovery

Excessive or prolonged activation of immune and stress responses can be detrimental to host tissues, ultimately compromising organismal survival (32, 33). The above-described results indicate that AIA neurons restrain innate immune and stress pathway activation during infection and suppress ER stress signaling during recovery. These findings raise the possibility that failure to appropriately regulate these responses results in tissue damage, and that AIA neuronal activity is required to preserve tissue integrity during both infection and recovery phases.

To test this hypothesis, we assessed intestinal integrity using two complementary approaches: the Smurf assay, a qualitative dye-leakage assay that reports gross intestinal barrier failure (37), and the propidium iodide (PI) assay, which provides a quantitative measure of epithelial damage at the cellular level (38, 39). Together, these assays allowed us to evaluate whether inactivation of AIA neurons exacerbates tissue damage during *S. enterica* infection and subsequent recovery.

The Smurf assay is a non-invasive method in which animals ingest a non-absorbable blue food dye that remains confined to the intestinal lumen in healthy animals but leaks into the body cavity when intestinal integrity is compromised (37). In *AIA::HisCl* animals without histamine treatment, dye localization was restricted to the intestine prior to pathogen exposure (Figure 4A). Following exposure to *S. enterica*, dye leakage became visible by 24 hours and progressively worsened at 48 hours, indicating infection-induced intestinal damage (Figure 4, B and C). Notably, silencing AIA neurons with histamine markedly increased both the frequency and severity of dye leakage at both time points, with widespread staining throughout the body cavity observed by 48 hours (Figure 4, B and C). These results demonstrate that AIA neuronal activity limits intestinal damage during active infection.

**Figure 4.**
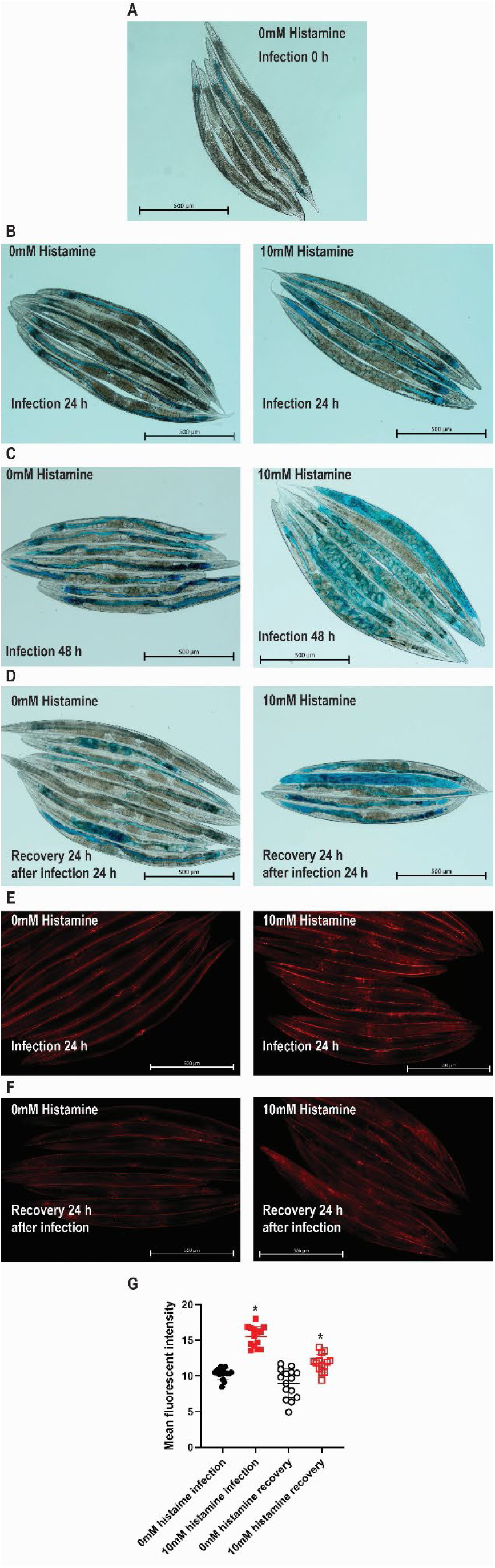
AIA neuronal activity is required to limit tissue damage during infection and recovery. Synchronized *AIA::HisCl* animals were exposed to *S. enterica* in the presence or absence of 10 mM histamine to selectively silence AIA neurons. **(A)** A representative image of animals prior to *S. enterica* exposure in the absence of histamine and soaked in blue food dye for 3 h to assess intestinal barrier integrity (Smurf assay). **(B)** Representative images of animals following 24 h exposure to *S. enterica* in the presence or absence of 10 mM histamine and subsequently soaked in blue food dye for 3 h to assess intestinal barrier integrity. **(C)** Representative images of animals following 48 h exposure to *S. enterica* in the presence or absence of 10 mM histamine and subsequently soaked in blue food dye for 3 h to assess intestinal barrier integrity. **(D)** Representative images of animals after 24 h recovery on *E. coli* OP50 in the presence or absence of 10 mM histamine following 24 h exposure to *S. enterica* and soaked in blue food dye to assess intestinal leakage. **(E)** Representative images of animals following 24 h exposure to *S. enterica* in the presence or absence of 10 mM histamine and soaked in propidium iodide (PI) for 3 h to visualize cellular damage. **(F)** Representative images of animals after 24 h recovery on *E. coli* OP50 in the presence or absence of 10 mM histamine following 24 h exposure to *S. enterica* and stained with PI to assess cellular damage. **(G)** Quantification of PI fluorescence intensity in animals treated with 0 mM or 10 mM histamine. Data are representative of three independent experiments (*n* = 15 animals per condition per experiment). Error bars represent SD. Statistical significance was determined using Sidak’s multiple comparisons test; Asterisks (*) denote *p* < 0.05 relative to the 0 mM histamine control.

We next examined intestinal integrity during the recovery phase using the Smurf assay. *AIA::HisCl* animals were exposed to *S. enterica* for 24 hours and then transferred to recovery plates containing *E. coli* OP50 and ampicillin to eliminate residual pathogen, in the presence or absence of histamine. Under recovery conditions without AIA silencing, dye leakage remained stable compared to 24 hours post infection (Figure 4D). In contrast, histamine-treated *AIA::HisCl* animals displayed pronounced dye leakage during recovery (Figure 4D), indicating worsening intestinal damage. These findings show that AIA neuronal activity is required not only to limit tissue damage during infection but also to promote restoration of tissue integrity during recovery.

We validated these findings using the PI assay. PI is excluded from intact intestinal epithelia but penetrates damaged cells, allowing quantification of tissue injury by PI fluorescence intensity. Consistent with the Smurf assay results, silencing AIA neurons significantly increased PI fluorescence in *AIA::HisCl* animals after 24 hours of *S. enterica* exposure, indicating enhanced intestinal damage during infection (Figure 4, E and G).

Similarly to the Smurf assays during recovery, histamine-treated *AIA::HisCl* animals exhibited significantly elevated PI fluorescence compared to controls without histamine treatment, demonstrating sustained epithelial damage when AIA neurons were inactivated (Figure 4, F and G).

Together, these results demonstrate that AIA neurons play a critical protective role in maintaining intestinal integrity during both infection and recovery. By constraining excessive immune and stress responses during infection and promoting their resolution during recovery, AIA neuronal activity prevents tissue damage and supports long-term host survival following pathogenic challenge.

## Discussion

In this study, we identify the AIA interneurons as a central regulator of immune homeostasis during both active infection and post-infection recovery in *C. elegans*. Using acute and chronic neuronal silencing, recovery-specific assays, and tissue damage assessments, we demonstrate that loss of AIA function results in sustained stress signaling, increased intestinal damage, and reduced survival. Our results suggest that AIA neuronal activity restrains excessive activation of innate immune and stress pathways during infection and is required for proper infection resolution by suppressing the XBP-1-mediated UPR during recovery. To our knowledge, this is the first study to identify a defined neuron that is specifically required for survival during the recovery phase following pathogen clearance.

Neuroimmune circuits in *C. elegans* are often described as immunosuppressive because their inactivation enhances immune gene expression and can transiently increase pathogen resistance (6). This raises a fundamental question: why would neural circuits dampen immune responses that are clearly required for host defense? Our findings clarify the biological logic underlying this regulation. During *S. enterica* infection, silencing AIA neurons resulted in excessive activation of PMK-1-dependent immune genes, insulin/IGF-1 pathway targets, and XBP-1-mediated ER stress genes (Figure 2). Despite heightened immune signaling, these animals exhibited reduced survival and increased intestinal damage (Figures 1 and 3), demonstrating that hyperactivation is detrimental. These observations are consistent with the broader principle that excessive inflammatory or stress signaling can drive tissue pathology (32, 33). Thus, the function of AIA neurons is not to maximize antimicrobial output, but to limit the amplitude of immune and stress responses and prevent collateral damage.

The importance of AIA activity becomes even more pronounced during recovery. Selective silencing of AIA neurons after pathogen clearance significantly reduced survival, and this defect was rescued by *xbp-1* RNAi (Figure 3), indicating that unresolved ER stress is the principal driver of post-infection mortality. Importantly, this finding demonstrates that resolution is not a passive decline of immune signaling but an actively regulated process that requires AIA neuronal activity. Collectively, our data support a model in which AIA-mediated neuroimmune regulation serves dual functions: constraining excessive immune and stress activation during infection and promoting timely suppression of ER stress during recovery.

Although multiple immune and stress pathways are activated during infection, sustained XBP-1-mediated UPR signaling is the dominant pathological factor during recovery when AIA function is lost, as the resultant survival defect was rescued by *xbp-1* RNAi but not by *pmk-1* or *daf-16* RNAi (Figure 3D). PMK-1 signaling, while essential for antimicrobial defense (7, 40), is not the major determinant of survival after pathogen clearance in this context. Chronic activation of ER stress pathways is increasingly recognized as a driver of inflammatory pathology and tissue dysfunction in mammals (20, 41). Consistent with this concept, our Smurf and PI assays showed that AIA silencing exacerbates intestinal barrier damage during both infection and recovery (Figure 4), linking persistent UPR activation to tissue injury. These findings suggest that neuronal control of ER stress resolution is essential for preserving tissue integrity following infection.

The recovery paradigm used here builds on the antibiotic-based infection-recovery model established by Aballay and colleagues (35, 36), who demonstrated that recovery requires active transcriptional reprogramming mediated by the intestinal GATA transcription factor ELT-2 and is associated with induction of detoxification and homeostatic pathways. Our findings extend this framework by identifying a specific neuron that is required for survival during recovery and by demonstrating that neuronal regulation of ER stress resolution is a key determinant of outcome. Together, these results support a model in which recovery involves coordinated transcriptional and neural programs that dampen stress responses and restore tissue homeostasis.

Given the evolutionary conservation of neuroimmune communication and UPR pathways (42, 43), similar neural mechanisms may regulate inflammation resolution in higher organisms. Indeed, failure of resolution is increasingly recognized as a central driver of chronic inflammatory diseases (21). By identifying a defined interneuron that actively controls infection recovery, our study provides mechanistic insight into how the nervous system coordinates not only host defense but also restoration of tissue homeostasis following pathogenic challenge.

## Materials and Methods

### Nematode Strains

The WT strain of *C. elegans* was Bristol *N2.* The neuron-specific histamine chloride channel (HisCl) strains were *CX15261(kyIs617)* [AIA expressing HisCl], *CX16863(kyIs698)* [AIY expressing HisCl], and *CX16781*(*kyIs617;kyIs698*) [AIA and AIY expressing HisCl]. Transgenic animals expressing GCaMP5a and HisCl in AIA neurons were generated by first injecting a *gcy-28d::GCaMP5a* transgene into WT animals, followed by injection of an *ins-1(short)p::HisCl1::SL2::mCherry* construct into the resulting strain. AIA-ablated strain *JN580(pels580)* was obtained from the *Caenorhabditis* Genetics Center (CGC, University of Minnesota, Minneapolis, MN). These strains were maintained as hermaphrodites at 20°C, grown on modified Nematode Growth Media (NGM) (0.35% instead of 0.25% peptone), and fed *E. coli* OP50 (44). All *C. elegans* animals used in this study were synchronized to 65-hours old unless specified otherwise.

### Bacterial strains

The following bacteria were grown using standard conditions: *Escherichia coli* OP50, *Escherichia coli* OP50::*gfp*, *Salmonella enterica* SL1344 (45).

### Survival assay

WT and HisCl animals were synchronized via egg-laying. Well-fed gravid adult animals were placed onto fresh *E. coli* OP50 NGM plates and incubated at 25°C for 45 minutes to lay eggs. After 45 minutes, the adult animals were removed from the plate and the eggs were allowed to grow for 65 hours at 20°C. *S. enterica* plates were prepared by culturing the bacteria for 15 - 16 hours in Luria broth (LB) in a shaking incubator at 37°C. A 30-µL drop of the fresh bacterial broth was placed on a 3.5cm NGM plate containing either 0 mM or 10 mM of histamine. Plates were incubated at 37°C for 15 - 16 hours then cooled to room temperature prior to use. Before *S. enterica* exposure, synchronized WT and HisCl animals were transferred to empty NGM plates containing 10 mM of histamine for at least 30 minutes to effectively silence the neurons (28, 29). Survival assays were carried out at 25°C and live animals were transferred daily until egg laying ceased. Animals were scored daily and were considered dead if they failed to respond to touch. Animals that crawl on the wall and died due to desiccation were censored from the data.

### Recovery assay

Synchronized animals were prepared as described above. Synchronized animals were transferred to fresh *S. enterica* plates containing 0 mM of histamine and incubated at 25°C for 24 hours. NGM plates containing 10 mM of histamine seeded with *E. coli* OP50::*gfp* were prepared using the method described above with the addition of 100 µg/mL ampicillin in the media. After 24 hours, animals were transferred to an empty NGM plate containing 10 mM of histamine for at least 30 minutes prior to being transferred to the *E. coli* OP50::*gfp* plates. Recovery assays were performed at 25°C and transferred daily until egg laying ceased. Animals were scored daily and were considered dead if they failed to respond to touch. Animals that crawl on the wall and died due to desiccation were censored from the data.

### Imaging of AIA neurons

#### Imaging setup and microfluidic devices

Calcium imaging was performed on a Leica DMI8 inverted microscope with a Leica DFC7000 camera. A Leica EL6000 light source was used for illumination. 12-bit images were acquired at 4 Hz with 170 msec exposure and 4x binning using a 40x/0.75 N.A. air objective.

Control of stimuli delivery, including odors with and without histamine, was done using microfluidic devices as previously described (46), with small modifications. Briefly, fast stimuli transitions (200-400 ms) were controlled using a flow-control switch within the microfluidic chamber, which was regulated by a Lee solenoid valve. ElveFlow twelve-way valves were located upstream of the microfluidic chamber input channels, allowing presentation of various stimuli to the animals. Flow dynamics was quantified using fluorescein in each experiment, allowing quality control and temporal alignment of imaging data.

The 30 μm tall microfluidic molds were fabricated as previously described (46) with small modifications. 4-inch wafers were baked at 250°C for 10 min. SU8-2025 was poured on the wafers and spun (10 sec at 450 rpm followed by 40 sec at 2250 rpm). Wafers were baked for 3 minutes at 65°C followed by 7 minutes at 95°C. Wafers were exposed to UV through the photomasks using an MA-6 mask aligner in low-vacuum contact mode (two 7 sec exposures separated by a 20 sec pause) and were then post-baked for 3 minutes at 95°C. The SU-8 layer was then developed for 2 minutes, transferred to fresh developer for 1 minute, and rinsed in IPA for 1 minute, followed by hard bake with a temperature ramp from 95°C up to 190°C over 80 minutes. The resulting molds were used to make ∼7 mm thick polydimethylsiloxane (PDMS) devices (Sylgard 184 A and B, mixed at 10:1 ratio).

#### Calcium imaging experiments

Fresh diacetyl odor stimuli (Sigma-Aldrich) diluted into the indicated final concentrations were prepared daily in S basal buffer in pre-cleaned amber glass vials (Environmental Sampling Supply). Imaging experiments were performed on one-day old adult hermaphrodite animals. Animals were washed with S basal buffer and placed on an NGM plate flooded with S basal containing the paralytic agent 4 mM (-)-Tetramisole hydrochloride (Sigma) for 10 minutes. Animals were then picked using a syringe, loaded into the microfluidic device, and monitored upon delivery of diacetyl at different concentrations. Each animal was exposed to six 30-second diacetyl pulses as follows: twice 115 nM, twice 1.15 μM, and twice 11.5 μM. Repeated presentations of the same stimulus concentration were separated by 30 sec, whereas different concentrations were separated by 60 sec.

For animals exposed to histamine, animals were grown overnight on NGM plates supplemented with 10 mM histamine. All subsequent washes and odor stimuli were performed using S basal buffer containing 10 mM histamine.

#### Image analysis of AIA imaging experiments

A MATLAB (MathWorks) graphical user interface (GUI) was developed to assign axonal compartments, followed by automated frame-to-frame ROI tracking and signal extraction. Raw signals were extracted by averaging the brightest fluorescent pixels within their assigned polygons, while the background fluorescence was calculated by averaging the dimmest fluorescent pixels in the surrounding area. Noise was filtered from background-subtracted signals using a 0.75-second moving average window, and neuronal activity (ΔF/F0) was calculated relative to the baseline fluorescence (F0), defined as the average fluorescence during the one-second preceding the increase in diacetyl concentration. Dye patterns were extracted from fluorescein by computing the mean fluorescence in an area near the animal nose. Dye data were used for quality assurance and to temporally align imaging data to the precise moment of stimulus arrival at the nose.

### RNA isolation

WT and HisCl gravid adult animals were lysed using a solution of sodium hydroxide and bleach (5:2). The freed eggs were washed with M9 buffer three times and allowed to hatch in S-basal liquid media at room temperature for 22 hours. Synchronized L1 larval animals were transferred to NGM plates seeded with *E. coli* OP50 and grown for 48 hours at 20°C until the animals reached the L4-larval stage. Animals were collected and washed in M9 buffer. Prior to transferring to *S. enterica* plates, animals were soaked in M9 buffer containing either 0 mM or 10 mM of histamine for 1 hour. After 1 hour, animals were transferred to NGM plates containing 0 mM or 10 mM of histamine and seeded with *S. enterica* for 4 hours at 25°C. After 4 hours, animals were collected and washed in M9 buffer. RNA was extracted using the NucleoZol and NucleoSpin RNA kit (Macherey-Nagel, Catalog No. 740406) following the protocol provided by the manufacturer.

### Quantitative real-time PCR (qRT-PCR)

Total RNA was obtained as described above. Five hundred ng of RNA was used to generate cDNA in a 20-µL reaction using the LunaScript RT super mix kit (New England Biolab, Catalog No. E3010). qRT-PCR was performed using the prescribed method for New England Biolab’s Luna Universal qPCR Master Mix on an Applied Biosystems StepOnePlus real-time PCR machine. Ten-µL reactions were set up following the manufacturer’s recommendations, and 10 ng of cDNA was used per reaction. Relative fold-change for each transcript was determined using the comparative C_T_(2^-ΔΔCT^) method and were normalized to pan-actin (*act-1*, *-3*, *-4*). Amplification cycle thresholds were determined by the StepOnePlus software. All samples were run in triplicate.

### RNA interference (RNAi)

RNAi was performed with the Ahringer group library by feeding *C. elegans E. coli* strain HT115 (DE3) expressing double-stranded RNA (dsRNA) that is homologous to the target gene for at least two generations. Before exposure, all RNAi clone plasmids were isolated and Sanger sequenced using a M13 reverse primer to confirm gene specificity. *E. coli* with the appropriate dsRNA vector were cultured in LB broth containing ampicillin (100 µg/mL) at 37°C for 15 - 16 hours and 120 µL was plated on NGM plates containing 100 µg/mL of ampicillin and 3 mM isopropyl-β-D-thiogalactoside (IPTG). The bacteria were allowed to grow for 15 - 16 hours at 37°C. The plates were cooled to room temperature in the absence of light before animals were transferred. Animals were synchronized using the method described above on the fresh RNAi plates. *unc-22* RNAi was included in each experiment as a positive control.

### SMURF assay

SMURF assays were done following the method described by Gelino et al.(37). Animals were synchronized as described above. Sixty-five-hour old *AIA::HisCl* animals were transferred to NGM plates containing 0 or 10mM of histamine seeded with *S. enterica* and incubated for 24 hours at 25°C. After 24 hours, animals were transferred to new NGM plates containing 0 or 10mM of histamine seeded with either *S. enterica* or *E. coli* and incubated at 25°C for another 24 hours. Animals were picked at specified time points and were transferred to 50 µL of a 5% solution of erioglaucine disodium salt in M9 buffer. The animals were allowed to soak in the erioglaucine solution for at least 3 hours before being washed with 1 mL of 30 mM sodium azide; the animals were centrifuged at 500 x *g* for 1 minute and the supernatant was discarded. Animals were washed 5 times to remove the erioglaucine solution then were mounted on a glass slide using a drop of halocarbon oil and imaged using a Zeiss Observer inverted microscope.

### Propidium iodide assay

*AIA::HisCl* animals were synchronized and exposed to *S. enterica* or *E. coli* in the presence of 0 or 10mM histamine as described above. Animals at specific time points were picked and transferred to 50 µL of 10 µM propidium iodide in M9 buffer and allowed to soak for at least 3 hours in the absence of light at room temperature. After 3 hours, the animals were washed 5 times with 30 mM sodium azide and centrifuged at 500 x *g* for 1 minute after each wash. The animals were then mounted to a glass slide using a drop of halocarbon oil and imaged using a Zeiss Observer inverted microscope.

### Quantification and statistical analysis

Survival curves were plotted using the Graphpad Prism computer software (version 10). Graphpad Prism uses the Kaplan-Meier method to calculate survival fractions and the log-rank (Mantel-Cox) test to compare survival curves. Survival curves of two experimental groups were considered significantly different when *p*-value < 0.05. qRT-PCR results were analyzed using a two-way ANOVA and Šídák’s multiple comparisons test to determine significance. All experiments were repeated at least three times, unless otherwise indicated. Statistical details for each figure are listed in the corresponding legend.

### Data, Materials, and Software Availability

The *C. elegans* strains and recombinant DNA generated in this study will be shared upon request, but we may require payment to cover shipment and completion of a Material Transfer Agreement for possible commercial applications. Any additional information required to reanalyze the data reported in this paper is available upon request.

## Acknowledgements

We thank Cori Bargmann for sharing reagents, Menachem Katz for assistance with microinjections, Amit Shacham (MNFU, Technion) for assistance with photolithography, and Nitsan Dahan (LSE, THHI, Technion) for assistance with microscopy. Some worm strains in this study were provided by the *Caenorhabditis* Genetics Center, which is funded by the NIH Office of Research Infrastructure Programs (P40 OD010440). This work was supported by NIH (R35GM124678 to J.S.) and The United States - Israel Binational Science Foundation (BSF2023287 to S.L. and J.S.). The funders had no role in study design, data collection and interpretation, or the decision to submit the work for publication.

## Author contributions

P.W., N.H., S.L., and J.S. designed experiments and analyzed data; P.W. and N.H. performed experiments; P.W., S.L., and J.S. wrote the paper.

## competing interests

The authors declare that they have no conflict of interest.

## Supporting Information

**Figure S1.**
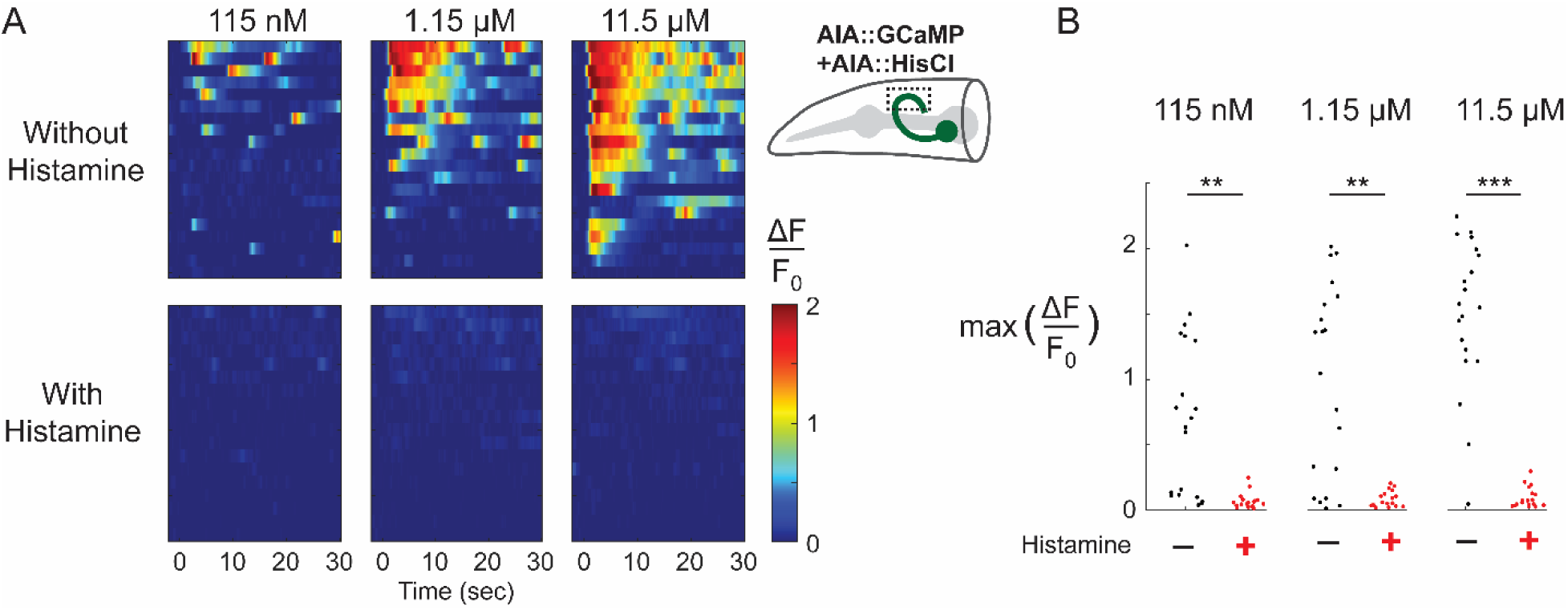
Chemogenetic silencing of AIA neurons by histamine-gated chloride channels (HisCl) in the presence of histamine. **(A)** AIA responses to diacetyl at the indicated concentrations in animals expressing HisCl in AIA neurons, in the absence (upper panels) or presence (lower panels) of histamine. Activity (ΔF/F_0_) was measured in the AIA axon (see schematic) and calculated relative to the baseline fluorescence (F₀) before odor delivery. Heat maps are sorted by the average response within the stimulus period. **(B)** Maximal AIA responses to diacetyl at the indicated concentrations in the absence (black) or presence (red) of histamine. *N* = 20 and *N* = 18 neuronal responses from the −histamine and +histamine conditions, respectively. Statistical significance was calculated using a Wilcoxon rank-sum test: ** *p* < 0.005, *** *p* < 10^-6^.

## Notes

### Competing Interest Statement

The authors have declared no competing interest.

### Summary of Updates

This revised version addresses the elevated basal expression of XBP-1-regulated unfolded protein response (UPR) genes observed in AIA::HisCl and AIA+AIY::HisCl animals in the absence of histamine. To determine whether basal HisCl activity causes unintended silencing of AIA neurons, we performed new calcium imaging experiments in animals expressing both GCaMP and HisCl in AIA neurons. These experiments showed that AIA neurons retain robust odor-evoked calcium responses in the absence of histamine, whereas histamine treatment efficiently suppresses neuronal activity, confirming effective histamine-dependent silencing. Although subtle basal effects of HisCl expression cannot be excluded, these findings argue against substantial AIA silencing in the absence of histamine and suggest that the elevated basal UPR gene expression is more likely attributable to strain-specific differences in gene expression. Accordingly, we have revised the Results to incorporate these findings. This version also changes the title to better reflect the content, includes a new supplementary figure (Figure S1) presenting the calcium imaging data, a new Imaging of AIA neurons section in the Materials and Methods, and the addition of a new co-author.

